# 3D segmentation of perivascular spaces on T1-weighted 3 Tesla MR images with a convolutional autoencoder and a U-shaped neural network

**DOI:** 10.1101/2020.11.25.397364

**Authors:** Philippe Boutinaud, Ami Tsuchida, Alexandre Laurent, Filipa Adonias, Zahra Hanifehlou, Victor Nozais, Violaine Verrecchia, Leonie Lampe, Junyi Zhang, Yi-Cheng Zhu, Christophe Tzourio, Bernard Mazoyer, Marc Joliot

## Abstract

We implemented a deep learning (DL) algorithm for the 3-dimensional segmentation of perivascular spaces (PVSs) in deep white matter (DWM) and basal ganglia (BG). This algorithm is based on an autoencoder and a U-shaped network (U-net), and was trained and tested using T1-weighted magnetic resonance imaging (MRI) data from a large database of 1,832 healthy young adults. An important feature of this approach is the ability to learn from relatively sparse data, which gives the present algorithm a major advantage over other DL algorithms. Here, we trained the algorithm with 40 T1-weighted MRI datasets in which all “visible” PVSs were manually annotated by an experienced operator. After learning, performance was assessed using another set of 10 MRI scans from the same database in which PVSs were also traced by the same operator and were checked by consensus with another experienced operator. The Sorensen-Dice coefficients for PVS voxel detection in DWM (resp. BG) were 0.51 (resp. 0.66), and 0.64 (resp. 0.71) for PVS cluster detection (volume threshold of 0.5 within a range of 0 to 1). Dice values above 0.90 could be reached for detecting PVSs larger than 10 mm^3^ and 0.95 for PVSs larger than 15 mm^3^. We then applied the trained algorithm to the rest of the database (1,782 individuals). The individual PVS load provided by the algorithm showed a high agreement with a semi-quantitative visual rating done by an independent expert rater, both for DWM and for BG. Finally, we applied the trained algorithm to an age-matched sample from another MRI database acquired using a different scanner. We obtained a very similar distribution of PVS load, demonstrating the interoperability of this algorithm.

## 1 Introduction

With increasing longevity, cognitive impairment, stroke, and dementia are currently major causes of disability and dependence in elderly individuals (Seshadri and Wolf, 2007), representing a huge societal burden. Cognitive impairment and dementia are mainly the result of a mix of vascular brain injury and neurodegeneration. Most often, vascular brain injury does not result in stroke, but manifest as covert small vessel disease (cSVD), a pathology highly prevalent in elderly individuals (Schmidt et al., 1999). Small vascular brain injuries of various types (white matter lesions, lacunes, microbleeds, and enlarged perivascular spaces) can be detected with brain magnetic resonance imaging (MRI; (Wardlaw et al., 2013)) in older community individuals. These lesions are associated with a risk of cognitive decline and dementia (review in (Debette et al., 2019)) and are now considered imaging markers of cSVD. Thus, there is considerable interest in assessing the burden of these lesions from an epidemiological point of view and for the early identification of individuals at risk of developing severe cognitive deterioration and/or dementia.

Among these lesions, perivascular spaces (PVSs; (Lawrence and Kubie, 1927)), also referred to as Virchow-Robin spaces, are of particular interest. PVSs are extracellular spaces containing interstitial fluid or cerebrospinal fluid surrounding cerebral small penetrating arteries and veins (Patek, 1941). PVSs are formed during development, accompanying brain angiogenesis (Marin-Padilla and Knopman, 2011). As such, they are physiological spaces that, when large enough, can be visible on brain MRI scans of healthy individuals (Maclullich et al., 2004; Yakushiji et al., 2014; Zhu et al., 2011). Animal models have shown evidence that both degenerative and vascular mechanisms can lead to enlarged PVSs (ePVSs), and several studies have shown that ePVS is associated with age and the risk of both cognitive deterioration (Passiak et al., 2019) and dementia (Francis et al., 2019; Zhu et al., 2010b); ePVS is also associated with the presence of other cSVD imaging markers in elderly individuals (Doubal et al., 2010; Potter et al., 2015; Ramirez et al., 2016; Zhu et al., 2010a). Overall, ePVS is now considered a hallmark and a very early anomaly of cSVD, so its assessment has recently become a major area of interest.

ePVS burden is commonly assessed on T1- or T2-weighted MR images using visual ratings with semi-quantitative rating scales (Adams et al., 2013; Potter et al., 2015; Zhu et al., 2011). However, such visual reading lacks precision and reproducibility, which limits its usability for longitudinal studies, and leads to overall loss of analytic power. Therefore, there is a strong need for unsupervised quantitative volumetric segmentation methods that could ideally identify every PVS in each individual as a 3-dimensional (3D) object in a perfectly reproducible manner. In fact, in the past few years, there have been several attempts at developing such automated PVS segmentation methods using broadly two different approaches, one based primarily on image processing (Ballerini et al., 2018; Boespflug et al., 2018; Gonzalez-Castro et al., 2017; Schwartz et al., 2019; Sepehrband et al., 2019; Wang et al., 2016; Zhang et al., 2017) and the other mainly based on deep learning (DL) (Dubost et al., 2020; Dubost et al., 2019; Jung et al., 2019; Lian et al., 2018; Sudre et al., 2019). The former approach is based on signal enhancement/noise reduction and/or specifically tailored morphological filters derived from the precise analysis of a few PVSs. The latter approach is based on a large set of convolutional filters (at different spatial resolutions) that extract features relevant for segmenting target objects, such as PVSs in the image. Such algorithms require a priori knowledge of the PVS number or locations on a subset of the input data. Both approaches have drawbacks: image processing methods are hampered by the large variance of PVS shapes and the signal-to-noise ratio (SNR), making it difficult to design a filter that will be optimal for the detection of all kinds of PVSs. Conversely, DL methods are sensitive to the quality and amount of a priori knowledge available: in particular, having a sufficiently large and reliable learning set of PVSs may be very difficult and cumbersome as it will require having one or several human operators manually tracing multiple PVSs on thousands of subjects. Both types of methods suffer from limited interoperability, as algorithms are usually tuned for the type of images they are designed or trained with.

Thus, despite several interesting attempts, there is still a need for an interoperable, fully unsupervised, and validated algorithm for the detection and quantification of PVSs in the entire brain volume. In the present study, we investigated the possibility of implementing such an approach using a class of DL methods based on autoencoders (Kingma and Welling, 2013) and U-shaped networks (U-nets, (Ronneberger et al., 2015)); a key feature of this approach is that the algorithm is able to learn from relatively sparse data, a major advantage over other DL algorithms. Others have used similar approaches for the PVS detection (Dubost et al., 2020; Lian et al., 2018). Dubost et al. (Dubost et al., 2020) presented a weakly supervised detection method based on U-net architecture that could be optimized with the PVS count, and applied it to a large dataset (2,200 subjects) of T2-weighted scans. Lian et al. (Lian et al., 2018) proposed a multi-channel (T2-weighted, enhanced T2-weighted, and probability map) fully convolutional network and applied it to a small set (20 subjects) of 3D patches of scans acquired at 7T. Here, we report a simpler U-net implementation, based on the T1-weighted (T1w) single-channel input, on a large database of 3D T1w whole-brain volumes acquired at 3T from 1,832 young adults. The learning set was composed of 40 T1w MRI scans from this database in which all “visible” PVSs were manually annotated by an experienced operator. After learning, algorithm performance was assessed using another set of 10 T1w MRI scans from the same database in which PVSs were also traced by the same operator. Next, we applied the algorithm to the rest of the MRi-Share database (1,782 individuals) and compared its output to a visual rating given by a trained rater based on a validated scale. Finally, we applied the algorithm to the T1w MRI images of age-matched subjects from the BIL&GIN database, acquired from a different scanner; subsequently, we compared the PVS distribution of both databases.

## 2 Methods

The brain MRI data were taken from the MRi-Share database (Tsuchida et al., 2020), a subcomponent of i-Share (internet-based Student Health Research enterprise, www.i-share.fr), a large prospective cohort study aiming to investigate French university student health. The MRi-Share database was designed to allow the investigation of structural and functional brain phenotypes in a sample of approximately 2,000 young adults in the post-adolescence period. In the present study, we included 1,832 i-Share participants who completed the full MRi-Share brain imaging examination and did not have any incidental findings on their brain MRI. The study sample age was 22.1 ± 2.3 years (mean ± SD, range: [18-35], median = 21.7 years), with a high proportion of women (72%, 1,320 women) as was the case with the rest of the i-Share cohort. The study was approved by the local ethics committee (Bordeaux, France).

For the purpose of the present study, we used the T1w scans acquired from each participant on the same Siemens Prisma 3-Tesla MRI scanner using a three-dimensional high-resolution MPRAGE sequence (TR =2000 ms; TE = 2.03 ms; flip angle = 8°; inversion time = 880 ms; field of view = 256 x 256 x 192 mm^3^; isotropic voxel size = 1×1×1 mm^3^, and in-plane acceleration = 2).

### 2.1 Manual segmentation and visual rating of PVS

#### 2.1.1 Manual segmentation of PVS in a subsample of 50 participants

Supratentorial PVSs are usually classified based on their location, either in the basal ganglia (BG) along lenticulostriate arteries or in the deep white matter (DWM) of the brain. These two types of PVS are usually rated separately and/or quantified since they were demonstrated to be differentially associated with SVD and dementia (Ding et al., 2017), as well as to have different genetic determinants (Duperron et al., 2018). Accordingly, to train and evaluate the performance of our detection algorithm, a subset of 50 individuals exhibiting varying amounts of visible PVS either in the DWM or in the BG were selected from the study sample by a neuroradiologist (BM) who reviewed the raw T1w images of the entire dataset. A trained investigator (AT) performed a voxelwise manual delineation of each PVS on the raw T1w images of each of these 50 individuals. Manual annotation of each PVS was performed using Medical Image Processing, Analysis and Visualization (MIPAV, v7.4.0). Specifically, each axial, coronal, and sagittal slice from the raw T1w scans from each individual was reviewed using the 3D view setting of MIPAV to detect PVSs in the DWM and BG regions. DWM PVSs are typically visible as tubular shapes, often running perpendicular to the cortical surface following the orientation of perforating vessels, whereas those in the BG are visible at the base of the basal ganglia along lenticulostriate arteries. Based on these shape and location characteristics, each visible PVS was segmented as best as possible using the MIPAV pen tool and occasionally expanded using the MIPAV paint grow tool that can automatically “paint” every neighboring voxel that has a lower intensity level than the selected voxel; the distance limitation set for the paint grow tool was 3 mm. The PVS segmented volume was saved as a binary mask in the individual native acquisition space. The PVS annotation procedure was first optimized by having the first ten MRI datasets reviewed by a second expert (LL) who recorded all potential disagreements she had, whether false positive or false negative according to her own opinion, for every dataset. These discordances were then jointly checked one by one by the two experts and were resolved by consensus between them. Subsequently, the remaining 40 MRI datasets were manually annotated for PVS by the first expert only.

#### 2.1.2 Visual rating of PVS burden in all participants

The global PVS burden estimated with our algorithm was also compared to a classical visual semi-quantitative assessment. For this, another investigator (JZ) visually rated the global PVS burden for each of the 1,832 individuals of the sample using a previously validated protocol and rating scale (Zhu et al., 2011). Briefly, for each individual, all axial slices of the T1w images were first examined to identify the slice containing the largest amount of PVS (one for DWM and one for the BG). The selected slice was then used to rate the burden of PVS by the number of spaces observed on a 4-level severity score as follows: for the BG, degree 1 when there were <5 PVSs, degree 2 when there were ≥5 and ≤10 PVSs, degree 3 when there were >10 PVSs but they were countable, and degree 4 when there were innumerable PVS; for cerebral DWM, degree 1 when there were <10 PVSS in the entire cerebral white matter, degree 2 when there were ≥10 PVSs in the total cerebral white matter but <10 in the slice with the largest number of PVSs, degree 3 when there were ≥10 and ≤20 PVSs in the slice with the most PVSs, and degree 4 when there were >20 PVSs in the slice with the most PVSs. The reliability of this visual rating was assessed by having the same investigator blindly rate two subsets of 60 individuals (one for the BG and one for DWM, with 40% of individuals common to both subsets) twice. The kappa concordance coefficients between the two ratings were 0.81 and 0.77 for DWM and the BG, respectively (both were significantly different from 0 at p < 10^-4^), and all discrepancies between the two ratings were minor, i.e., consisting of a difference of one scale level.

### 2.2 Segmentation model training, validation and testing

This section details the methodology for the PVS segmentation model architecture, training, and testing. KNIME 4.0 (Berthold et al., 2009) was used for the data management workflows, and Python-based Keras 2.2.4 (https://keras.io), Scikit-learn (Pedregosa et al., 2011) and TensorFlow 2.1 (Abadi et al., 2016) were used for implementing our segmentation model. The algorithms were run on a Centos computer with a Xeon ES2640, 40 cores, 256 Gb RAM and two Tesla P100 GPUs with 16 Gb RAM.

#### 2.2.1 Methodology of PVS segmentation

##### 2.2.1.1 Defining data subsets for training and testing the PVS segmentation algorithm

The full dataset of 1,832 T1w volumes was split into 3 non-overlapping subsets:

- An autoencoder subset (ENCOD), including the 1,782 volumes without manual annotations of PVSs; the ENCOD subset was used to train the 3D convolutional autoencoder (see below), as well as to test the model-predicted PVS load against visual rating.
- A training subset (TRAIN), including 40 of the volumes with manually annotated PVSs; the TRAIN subset was used to train and validate the segmentation model.
- A testing subset (EVAL), including the remaining 10 volumes with annotated PVS; the EVAL subset was used to test the segmentation model.

The 10 T1w volumes of the EVAL subset were selected by an expert neuroradiologist (BM) to be representative of the full set of 50 volumes of manually annotated PVSs. This was checked by comparing the distribution of manually annotated PVSs in the two subsets.

##### 2.2.1.2 T1-weighted MRI preprocessing

Prior to training the DL, T1w MRI volumes (N=1,832) were preprocessed following a 5-step procedure: 1-tissue segmentation with FreeSurfer, 2-creation of an intracranial volume mask (ICV), 3-voxel intensity rescaling, 4-creation of a brain volume bounding box, and 5-creation of a BG mask.

- First, each T1w volume was segmented using FreeSurfer (v6.0, https://surfer.nmr.mgh.harvard.edu/), and the different tissue components were identified.
- Second, an ICV mask was defined as the union of the gray matter (GM), WM, and cerebrospinal fluid (CSF) tissue voxels.
- Third, voxel intensity values were linearly rescaled between 0 and 1 by setting the 99th percentile of each subject’s sample as the maximum. The values greater than 1 were set back to 1.
- Fourth, for each individual, we computed the minimal bounding box (oriented along the 3 axes of the T1w acquisition) that included his/her brain volume. In doing so, we eliminated the neck and some of the background air signals. The union of the 1,832 individual bounding boxes (registered using their centers) was then computed and used to crop each T1w volume. Note that in the process, T1w volumes were not interpolated but were only translated by an integer number of voxels since all individual boxes had the same orientation. This cropping process led to a 52% data size reduction (from 256×256×192 voxels to 160×214×176 voxels), resulting in a gain of a factor of approximately 2 in computational burden.
- Fifth, since PVS distribution is usually separately examined when localized in the BG or in the DWM, we created a basal ganglia mask (BG-mask) for each individual including the tissue classes identified by FreeSurfer as thalamus, thalamus-proper, caudate, putamen, pallidum and accumbens regions. The labels were used to determine whether a PVS belonged to the BG or the DWM.

##### 2.2.1.3 Autoencoder and U-net architecture

Our segmentation model used a U-net architecture similar to the one described in (Ronneberger et al., 2015). The main constraints in our application were (1) the small size of the annotated datasets available for training and validation (TRAIN set) and (2) the large volume of data: the model parameters and a batch of volumes had to fit into the 16 Gb memory of the GPU.

To train the U-net, an autoencoder was first trained on the large ENCOD set. The convolutional autoencoder and the U-net share a similar architecture to transfer the weights learned by the former to the latter. The difference resides in the addition of “skip connections” in the U-net between the corresponding encoding and decoding blocks (Figure 1); the output of the encoding block is concatenated to the input of the decoding block. We used a U-net architecture denoted by its main hyperparameters: the initial number of kernels for the first stage of convolutions (*nb_kernel_init = 8*), the number of stages (NStages = 7,) and the number of 3D convolutions for a stage (*nConvolutions = 2*). This configuration is referred in the following as the 8.7.2 autoencoder/U-net architecture.

**Figure 1:**
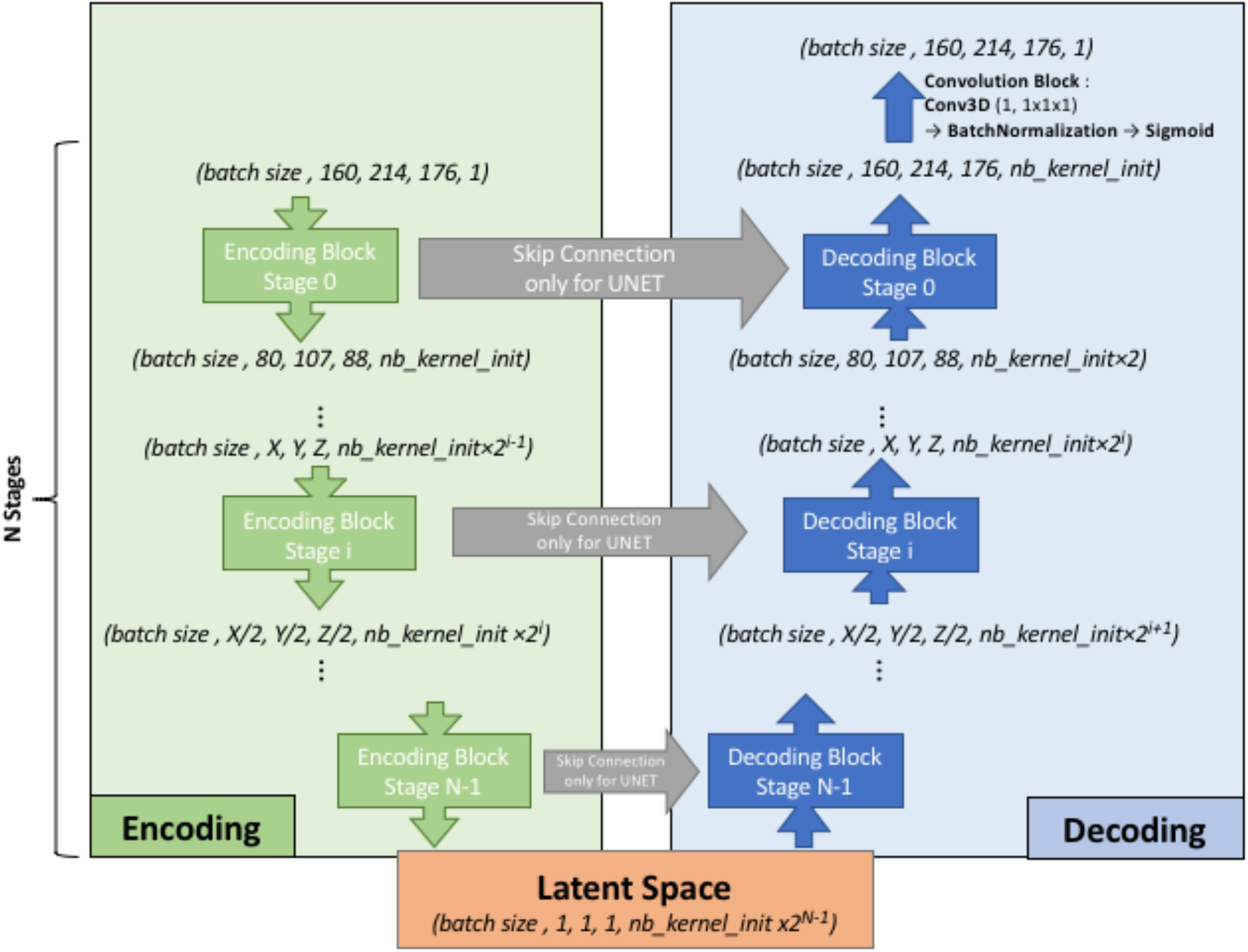
Overall architecture for the convolutional autoencoder (without skip connections) and U-net (with skip connections). On the left, the encoding blocks are shown. Each block used 3D convolutions to create new features and max pooling operations to downsize the image. On the right are the decoding blocks that used upsampling and convolutions to reconstruct the image (for the autoencoder configuration) or to create the segmentation probability map (for U-net configuration with skip connections).

Figure 2 shows the encoding and decoding levels for stage *i* of the encoding/decoding architecture. On the encoding side, the tensors went through 2 convolution blocks. Each convolution block consisted a 3D convolution with a kernel size of 3×3×3 that produced 2^i^ kernels, followed by batch normalization (Ioffe and Szegedy, 2015), and activation using a rectified linear unit (ReLU) (Glorot and Bengio, 2010). In the autoencoder, the weights were initialized randomly using a Glorot uniform initializer (Glorot and Bengio, 2010); in the U-net the weights were initialized using the trained autoencoder. ReLUs were used for all activation functions except for the last step of the decoding block, where a sigmoid was used to return to the initial volume or mask domain. After the convolution blocks, there was a 2×2×2 max pooling (Ciresan et al., 2011) and a dropout layer (Srivastava et al., 2014). We tested an architecture with strided convolutions instead of max pooling layers (as in (Milletari et al., 2016)), but the added parameters produced a model that did not fit into the GPU without reducing the number of kernels used at each stage, and the resulting model did not perform better.

**Figure 2:**
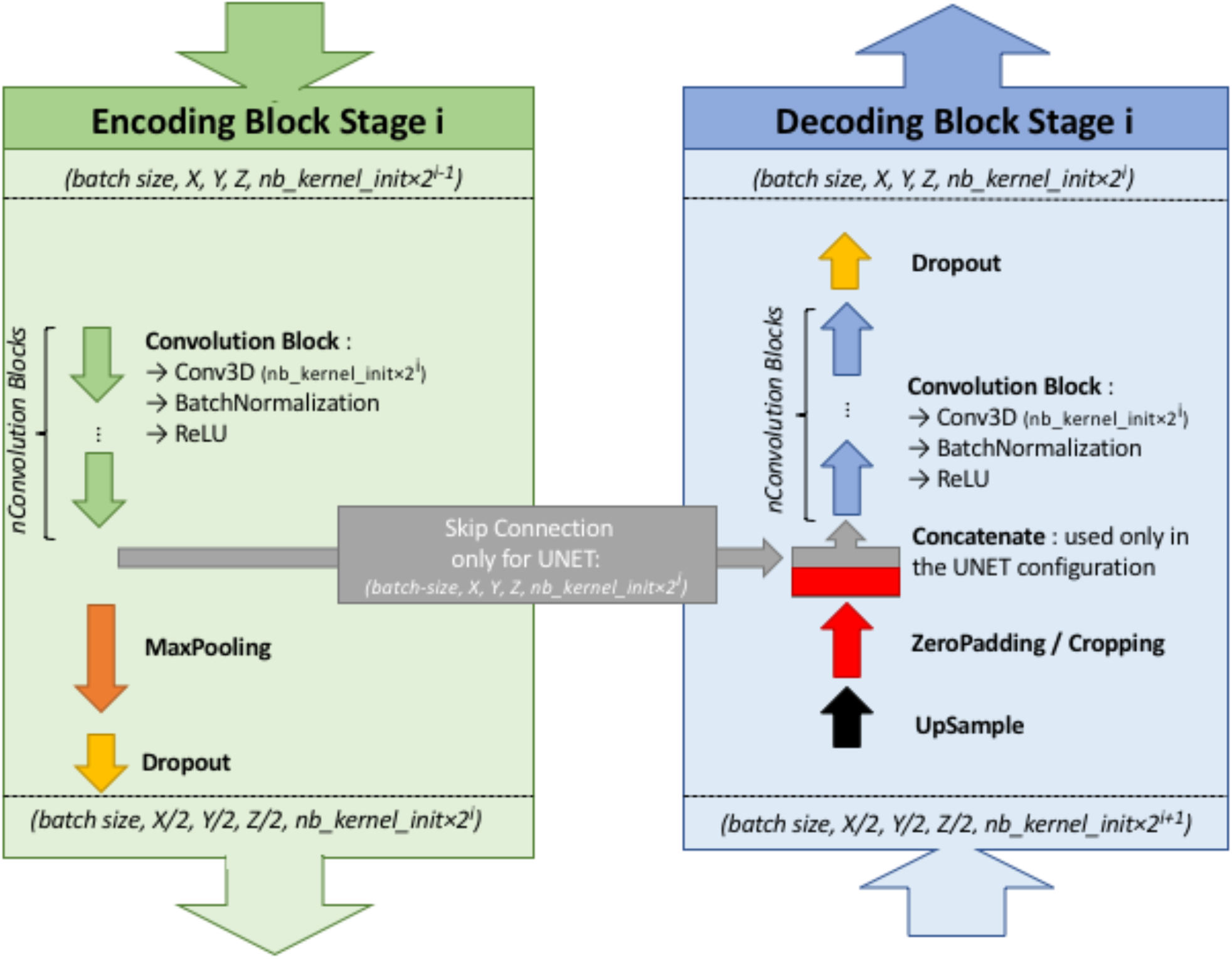
Details of stage i of the model architecture

On the decoding side, the first layer was a 2×2×2 nearest neighbor upsampling. Then, a padding/cropping layer was used to align the spatial dimensions of the tensor to those in the encoding block at the same stage. It allowed the subsequent concatenation of the tensors coming from the encoding block to those from the decoding layers in the U-net configuration, and eliminated the need for initial padding or cropping when the volume had resolutions that were not a power of 2. It was followed by a symmetric number of convolution blocks as the encoding side (i.e. 2 in our case), and finally, by a dropout layer. Unlike the encoding convolutions, the decoding convolutions were initialized using a Glorot uniform initializer for both the autoencoder and the U-net.

With our configuration of the 8.7.2 autoencoder/U-net architecture, the final model had 44 million trainable parameters. The resulting latent space had 512 dimensions. The Adam algorithm (Kingma and Welling, 2013) was used with the default parameters recommended by the authors: beta1=0.9, beta2=0.999, and decay=0.0. The dropout rate used for both encoding and decoding dropout layers was 0.1. In the autoencoder, the mean square error (MSE) was used as the loss function, whereas Keras implementation of the Dice loss was used as the loss function in the U-net. As we did not have enough computing resources to systematically explore the hyperparameter space for each possible configuration, the different hyperparameter values were chosen based on previous experiences. The most sensitive hyperparameter was the initial kernel size (larger was better), and we tried several configurations to balance it with the batch size while allowing the model and the batch of volumes to fit in GPU memory, resulting in a batch size of 2.

#### 2.2.2 PVS segmentation model training and testing

We first trained the autoencoder by randomly splitting the ENCOD dataset into 80% training and 20% validation subsets. We then initialized the U-net segmentation model with the weights of the first encoding stages and trained it using the TRAIN set with a 5-fold cross-validation scheme. Each fold consisted of 80% training (32 volumes) and 20% (8 volumes) validation, and each subject’s data appeared only once in the fold validation set and four times in the fold training set. The repartition in the 5-fold validation set was manually defined to include a similar pseudo-uniform distribution of the individual PVS load in each fold. For each fold, the model with the lowest validation loss was selected. The five resulting models were used to predict PVS maps for the EVAL subset, and the five output maps for each input image were averaged to create the final PVS map called thereafter the *consensus* (of the 5 folds) segmentation. Note that each predicted map was coded in values in the 0 to 1 range. Four indexes were computed to quantify segmentation algorithm performance on N=8 validation set for each of the 5 folds of TRAIN and N=10 of the EVAL sets. For this quantification, the prediction maps were binarized with a 0.5 threshold and compared to the manually traced PVS masks using *Dice-loss* scores (*see 2.2.1.3 above*.), the *true positive rate* (TPR) and the *positive predictive value* (PPV), as well as their harmonic mean known as the *(Sorensen-) Dice coefficient*. The *TPR* is defined as the number of predicted PVS voxels that were correctly identified (true positive, TP), i.e., overlapping with manually annotated PVS voxels, divided by the number of manually annotated PVS voxels (i.e. TP plus false negative (FN) voxels). The *PPV* is defined as the number of TPs divided by the sum of the number of TPs plus the number of false positive (FP) voxels, i.e., voxels that were wrongly predicted as PVSs. Note that in the context of machine learning, the *TPR* and *PPV* are also named *sensitivity* and *precision*, respectively. The Sorensen-Dice coefficient is then computed as 1/Dice = (1/TPR + 1/PPV)/2. The 3 indexes will be referred to as voxel-level indexes (VL) in the following sections.

In the testing phase, only the EVAL consensus prediction was used, and VL indexes were computed with 9 amplitude thresholds applied to the prediction map (*PredMap-Thr*, 0.1 to 0.9 in 0.1 step). Additionally, cluster-level (CL) TPR and PPV were computed as following: Clusters were defined using a 26-neighbor connectivity rule (i.e., 1 voxel is connected to its 26 surrounding voxels) both for the manually annotated and for the predicted PVS masks. A cluster of the predicted PVS volume was assigned to the TP class if it included at least one voxel of the manually annotated PVS volume. Similarly, a cluster of the predicted PVS volume was assigned to the FP class if it included no voxels of the manually annotated PVS volume. Conversely, a cluster of the manually annotated PVS volume was assigned to the FN class if it included no voxels of the predicted PVS volume. We will refer them as TPR-CL and PPV-CL, whereas their harmonic mean will be referred as Dice-CL. The final metrics were computed by averaging the values of the 10 EVAL subjects for each *PredMap-Thr* and each index. Note that the EVAL dataset (10 volumes) was not used in the training/validation step; thus, independent scores were generated.

### 2.3 PVS segmentation model robustness

In order to test the reproducibility of the model, we computed the segmentation model 4 more times. In addition, we assessed the robustness of the model with regard to the size of the training dataset by training the model with the reduced training set of 20 or 30 volumes. In each case, we present graphs of VL and CL TPR vs PPV for the 9 *PredMap-Thr*.

### 2.4 Testing the PVS segmentation model against the manual annotation: EVAL subset

This section reports the results obtained using the first segmentation model (based on 40 TRAIN participants’ data) with the EVAL subset.

#### 2.4.1 PVS segmentation model performance

Algorithm performance was assessed in the BG or in the DWM defined individually (see 2.2.1.). For each tissue type and for each subject of the EVAL set, the 4 quantification indexes (TPR-VL, PPV-VL, TPR-CL, and PPV-CL) and their harmonic means (Dice-VL and Dice-CL) (see 1.3.2) were computed independently for each of the 9 *PredMap-Thr* (0.1 to 0.9 in 0.1 step).

#### 2.4.2 PVS segmentation model performance for varying PVS cluster sizes

Since enlarged PVS is of primary interest, we investigated how TPR-CL and PPV-CL were modified if we focus on PVSs that were larger than a given threshold. Accordingly, TPR-CL vs PPV-CL plots with the 9 *PredMap-Thr* were generated for cluster size thresholds varying from 0 to 15 voxels with a step size of one voxel. For this analysis, the index values were computed using all the clusters of the EVAL set, not the average value for each subject.

To compute the TPR-CL, in this approach, we removed all clusters with a size smaller than a given threshold from the manually annotated map. Accordingly, TPs were clusters of the predicted map that had at least one voxel in common with a cluster of the thresholded annotated map. Meanwhile, to compute the PPV-CL, we removed the clusters smaller than a given threshold in the predicted map and compared that map with the annotated map.

#### 2.4.3 PVS segmentation model performance at predicting PVS cluster sizes

Finally, to assess the ability of the model to estimate the PVS size, we compared the size of manually annotated PVS clusters to that of the model-predicted clusters (i.e. TP clusters). For each cluster in DWM or BG, we computed the linear fit with the model-predicted cluster size as the independent and manually annotated cluster size as the dependent variables, applying 3 of the *PredMap-Thrs* (0.1, 0.5 and 0.9). We report both the slope and the R^2^ of the fit. Note that predicted clusters encompassed more than one manually traced cluster were not considered in this analysis. In percentage of the total number of TP clusters, the number of clusters removed was 7.6%, 3.6% and 2% for *PredMap-Thrs* of 0.1, 0.5, and 0.9, respectively.

### 2.5 Testing the PVS segmentation model of a large subset: ENCOD subset analysis

We used the first segmentation model (trained on 40 TRAIN participants’ data) to estimate the PVS load in the larger ENCOD subsets, in which the visual rating of PVSs was available for both DWM and BG. For each participant, we computed both the numbers of PVS voxels and of PVS clusters at three values of the *PredMap-Thrs* (0.1, 0.5, 0.9). We examined the relationship between the two numbers, searching for the best polynomial fit (up to 3^rd^ degree).

We then used the number of clusters as the proxy for the PVS load and compared it to the visual rating score, since the visual rating was based primarily on the evaluation of a number of PVSs. A logistic regression analysis was used to predict the visual rating score from the PVS cluster number for the whole, ENCOD set individuals (N=1,782), separately for DWM and for BG.

### 2.6 Assessment of the prediction algorithm interoperability

To indirectly assess the interoperability of our model, we predicted the PVS load in the T1w images from the BIL&GIN database (Mazoyer et al., 2015) that were acquired using a different MRI scanner (Philips Achieva 3T) 10 years prior to the acquisition of the data from the MRi-Share cohort, using the segmentation model trained on the MRi-Share dataset. T1w images were acquired using a different sequence (3D-FFE-TFE; TR = 20 ms, TE = 4.6 ms, flip angle= 10°, inversion time = 800 ms, turbo field echo factor = 65, sense factor = 2, matrix size = 256 x 256 x 180 mm^3^, and 1 mm^3^ isotropic voxel size). From the 453 subjects included in the BIL&GIN database, 354 were selected for being aged between 18 and 35 years to match the age range in the MRi-Share cohort. We assessed the similarity of PVS distributions in these age-matched cohorts using a QQ plot and tested using the Kolmogorov-Smirnov test. The tests were also computed using cluster size thresholds varying from 0 to 15 voxels by a step size of one voxel, and distributions were compared using the Kolmogorov-Smirnov test.

## 3 Results

### 3.1 Manual segmentation of PVS

The distribution of the number of voxels manually identified as DWM PVS (Figure 3.A) in 50 participants showed a bimodal shape with 34 participants below 1500 (1.5 cm^3^), 14 participants between 1500 and 3000 (1.5 and 3.0 cm^3^) and 2 outliers above 4000 (4 cm^3^). The distribution of the number of clusters in the DWM of the same participants (Figure 3.C) shows the same shape than the voxel distribution. For the PVS in the BG (Figure 3.B), the distribution was more homogeneous, with an average of 180 voxels (or mm^3^) and 2 outliers above 500 voxels (or mm^3^). The distribution of the number of clusters (Figure 3.D) had the same shape as the voxel distributions. Figure 3.E shows representative PVS of both categories. On average, in the whole set, a PVS in the BG was 2 times larger than a PVS in DWM.

**Figure 3:**
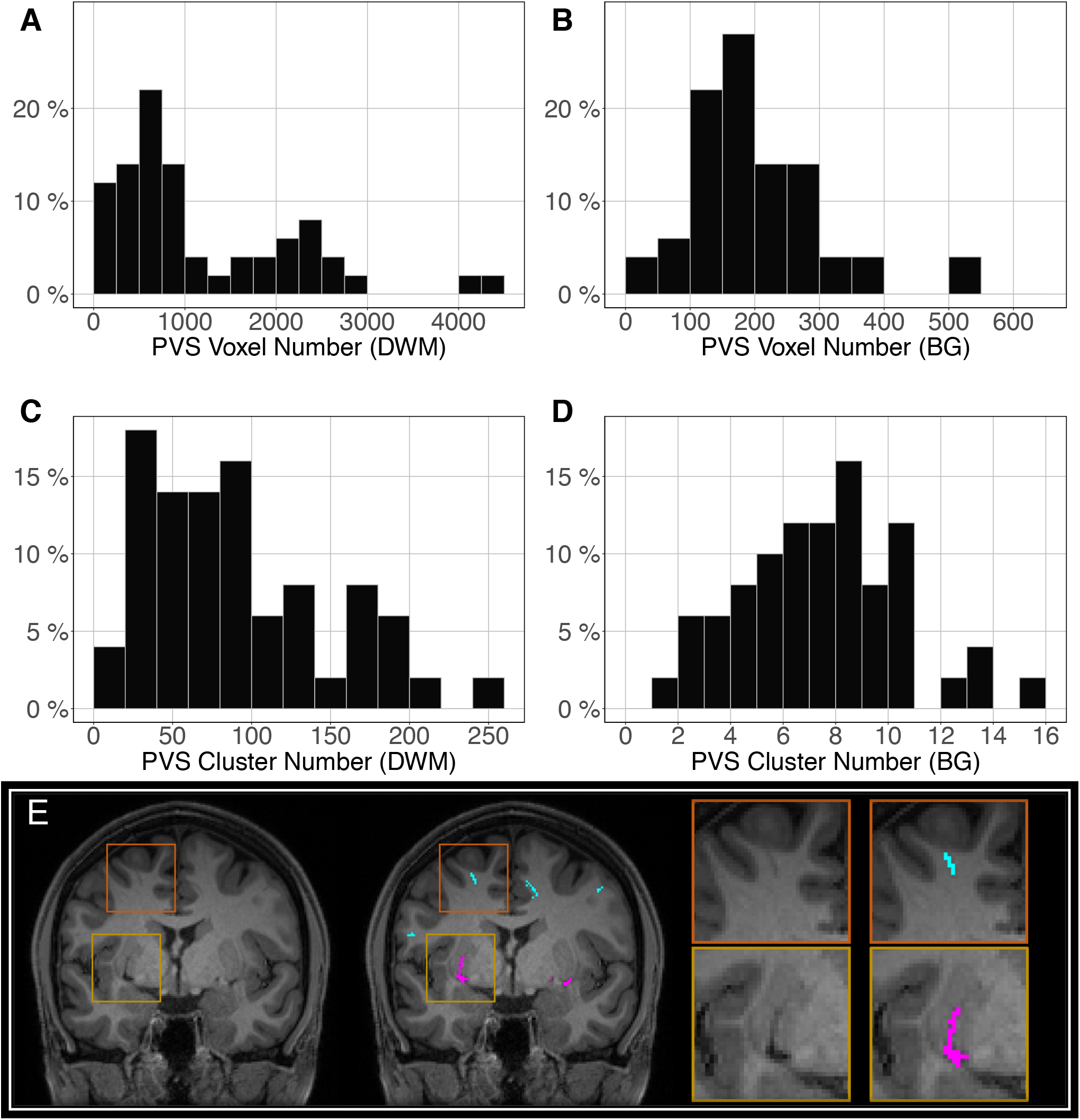
Distribution of the manually delineated PVS voxel load in the deep white matter (A) and basal ganglia (B) of 50 participants. Distribution of the manually delineated PVS number of clusters in the deep white matter (C) and basal ganglia (D) of 50 participants. (E) Example of delineated PVSs displayed over an MRI slice in one participant in the deep white matter (cyan) and basal ganglia (magenta). DWM: deep white matter, BG: basal ganglia.

### 3.2 Segmentation model training and testing

The most time-consuming step was the autoencoder training using the ENCOD dataset; one epoch run required ~2000 seconds for 1,425 volumes with a validation subset of 357 volumes. Figure 4.A shows the evolution of the loss function while training the autoencoder using the ENCOD dataset. The training was stopped after 23 epochs, and previous experiments showed that further training the autoencoder do not improved significantly the training speed of the *U-net*. Figure 4.B shows the loss evolution on the TRAIN dataset with 80/20 split training/validation for one-fold cross-validation. In this fold, the model with the lowest validation loss (0.4894) at epoch 119 was selected. Table 1 synthesized the 6 scores of each fold, computed as the average of one-fifth (8) of the 40 volumes of the TRAIN sets. Table 2 shows the average of the 10 EVAL volume result scores for each fold and for the consensus segmentation of the full model.

**Figure 4:**
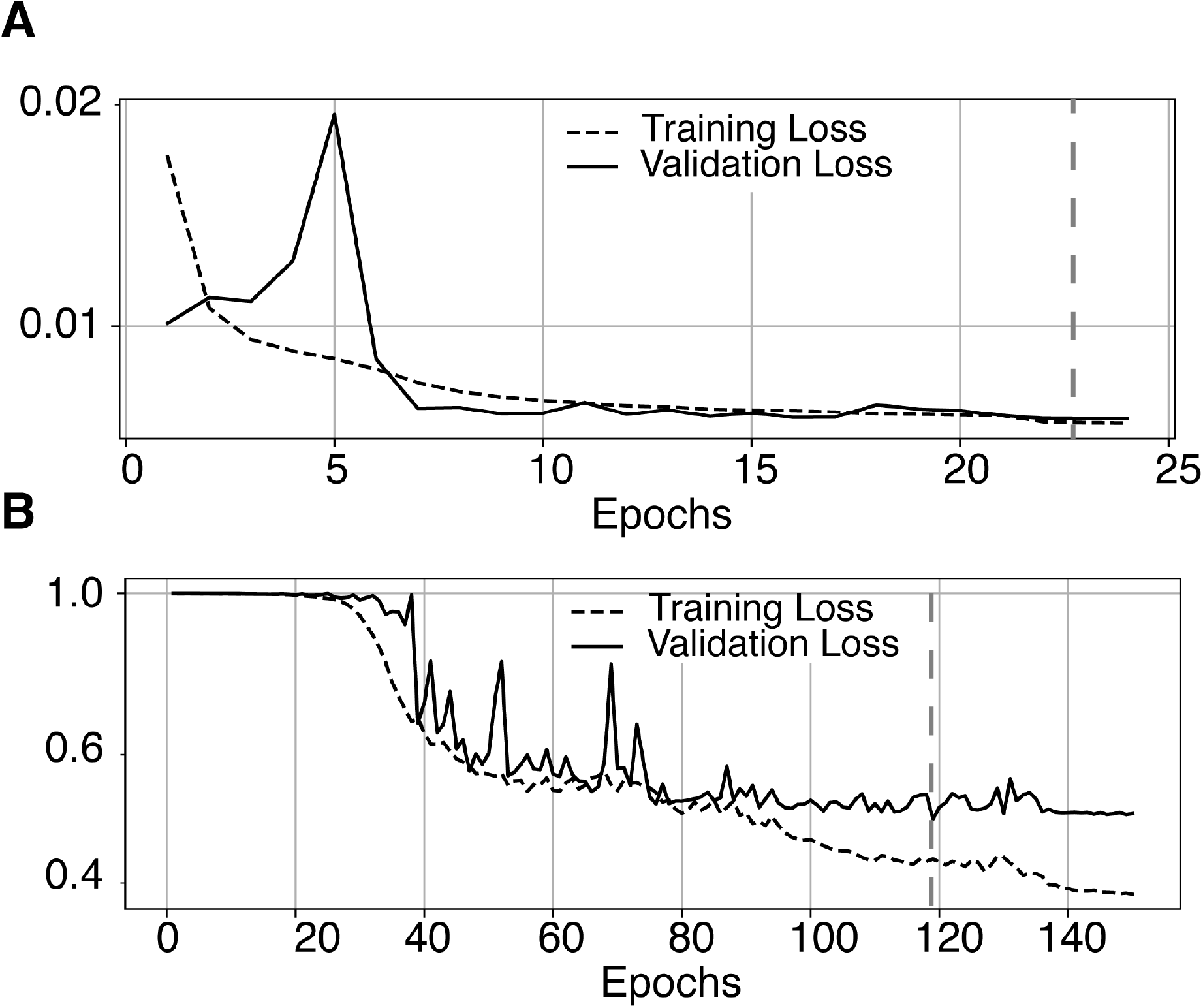
A) Training loss (MSE, dashed line) and validation loss (continuous line) evolution for the 8.7.2 autoencoder training. The model with the lowest validation loss (0.0075, epoch 23, dashed line) was selected. B) One-fold training loss (dice loss, dashed line) and validation loss (continuous line) evolution of 8.7.2. U-net with weights initialized from the autoencoder. The model with the best validation loss (0.49, epoch 119, dashed line) was selected. Logarithm scale is used for loss.

**Table 1:** Dice loss, true positive rate (TPR-VL), positive predictive value (PPV-VL) and Dice (Dice-VL) scores for each fold of the validation part of the TRAIN dataset (8 volumes). TPR: true positive rate, PPV: positive predictive value, VL: voxel level, CL: cluster level.

| Fold | Dice-loss | TPR-VL | PPV-VL | Dice-VL |
| --- | --- | --- | --- | --- |
| 0 | 0.483 | 0.523 | 0.556 | 0.539 |
| 1 | 0.513^ | 0.492 | 0.570 | 0.528 |
| 2 | 0.496 | 0.463^ | 0.635* | 0.535 |
| 3 | 0.501 | 0.510 | 0.521^ | 0.516^ |
| 4 | 0.471* | 0.524* | 0.578 | 0.550* |

\*: Best values of the 5 folds, ^: Worst values of the 5 folds

**Table 2:** Dice loss, true positive rate (TPR-VL), positive predictive value (PPV-VL) and Dice (Dice-VL) scores for each fold and for the 5-fold average of the EVAL dataset (10 volumes). TPR: true positive rate, PP: positive predictive value, V: voxel level, CL: cluster level.

| Fold | Dice-loss | TPR-VL | PPV-VL | Dice-VL |
| --- | --- | --- | --- | --- |
| 0 | 0.452 | 0.526 | 0.578 | 0.551 |
| 1 | 0.445 | 0.541 | 0.579 | 0.559 |
| 2 | 0.438* | 0.509 | 0.637* | 0.566* |
| 3 | 0.466^ | 0.561* | 0.520^ | 0.540^ |
| 4 | 0.443 | 0.503^ | 0.633 | 0.561 |
| 5-fold average | 0.573 | 0.526 | 0.634 | 0.575 |

\*: Best values of the 5 folds, ^: Worst values of the 5 folds

### 3.3 PVS segmentation model robustness analysis

Figure 5.A illustrates the good reproducibility of the model predictions across the 5 repetitions. Figure 5.B shows that even when the TRAIN dataset size was reduced to 30 individuals, performance was only slightly degraded compared to that obtained with the 40-subject training. While trying to train the algorithm using 20 datasets only, the algorithm performance visibly deteriorated, especially when analyzing the lowest *PredMap-Thrs* (between 0.1 and 0.4). Qualitatively, the degraded performance manifested as some predicted PVS clusters encompassing a large portion of the WM in some subjects of the TRAIN dataset, producing numerous FP voxels.

**Figure 5:**
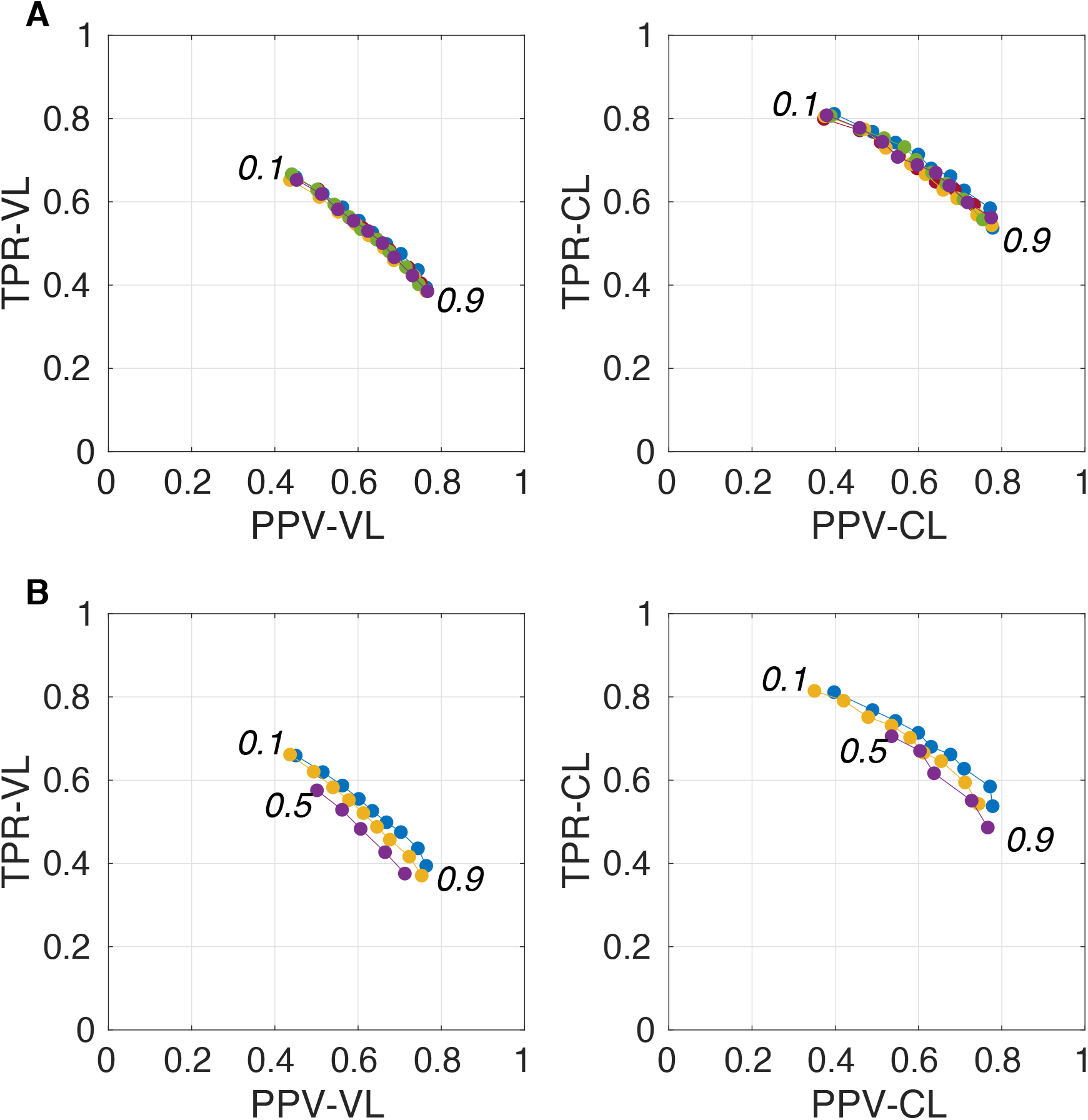
Plot of true positive rates (TPRs) versus positive predictive values (PPVs) for the 9-amplitude prediction map thresholds (PredMap-Thrs, 0.1 to 0.9 in step sizes of 0.1) computed for replication analysis based on 40 TRAIN subjects (each of the 5 replications is shown with a different color) (A) and for training the model with 20 (purple), 30 (yellow) and 40 (blue) subjects (B) using voxel level (VL) (left) and cluster level (CL) (right).

### 3.4 Testing the PVS segmentation model: EVAL-based subset analysis

#### 3.4.1 PVS segmentation model performance

Figure 6 shows for one subject of the EVAL subset, the visual display centered on one of the clusters identified as a TP. Such pictures were computed for each of the TP/FP/FN clusters.

**Figure 6.**
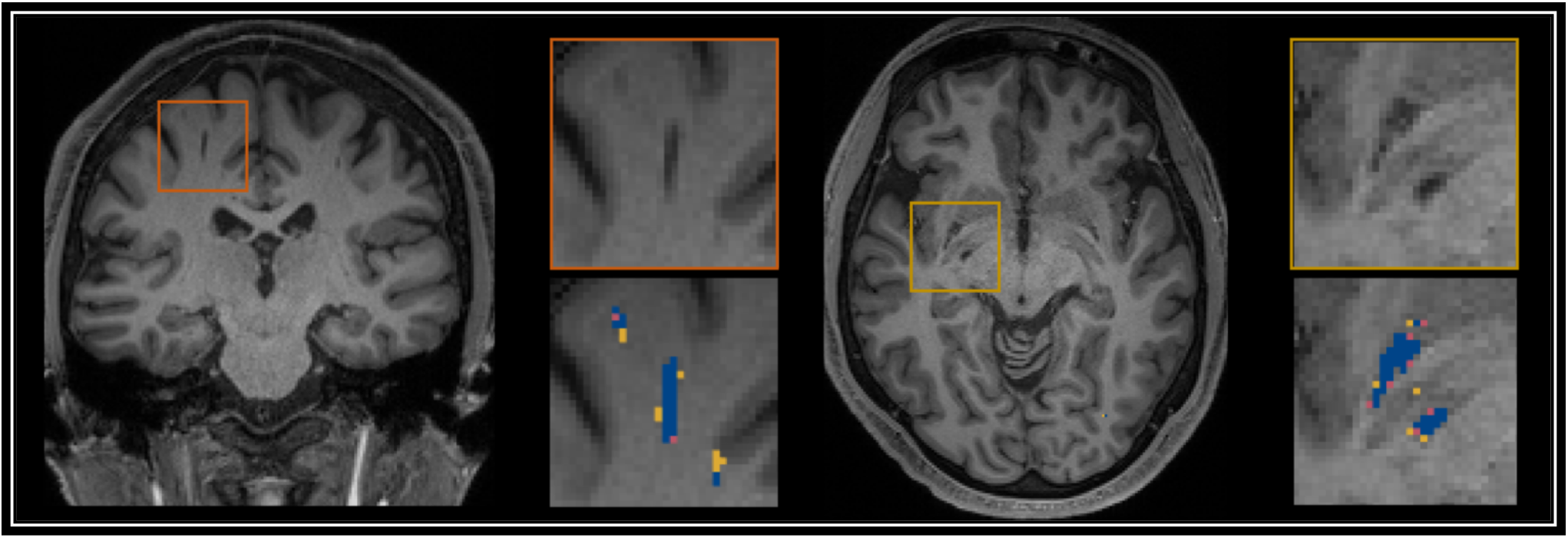
Example of true positive clusters detected and displayed in coronal and axial orientations. Blue indicates TP voxels, orange indicates FP voxels, and yellow indicates FN voxels.

Figure 7 shows plots of TPRs vs PPVs of segmented PVSs for the 9 *PredMap-Thrs* at both the voxel and cluster levels. Table 3 and Table 4 show the index values with Dice values. As expected, TPRs decreased and PPVs increased with increasing *PredMap-Thrs*, in both DWM and the BG, and at both VL and CL. While the PPV was similar at both VL and CL, the TPR was markedly higher at the cluster level. Regardless of the type of tissue (DWM/BG) and the level (VL/CL), the Dice coefficients were maximal for *PredMap-Thrs* between 0.4 and 0.6. This is unsurprising, considering that Dice coefficient is the harmonic mean of the TPR and PPV. More precisely, the best Dice values were for an amplitude of 0.6 in the DWM and for an amplitude of 0.4 in the BG. When comparing Dice coefficients on both levels, the impact of the prediction map amplitude thresholding seemed to be higher at the CL than at the VL.

**Figure 7:**
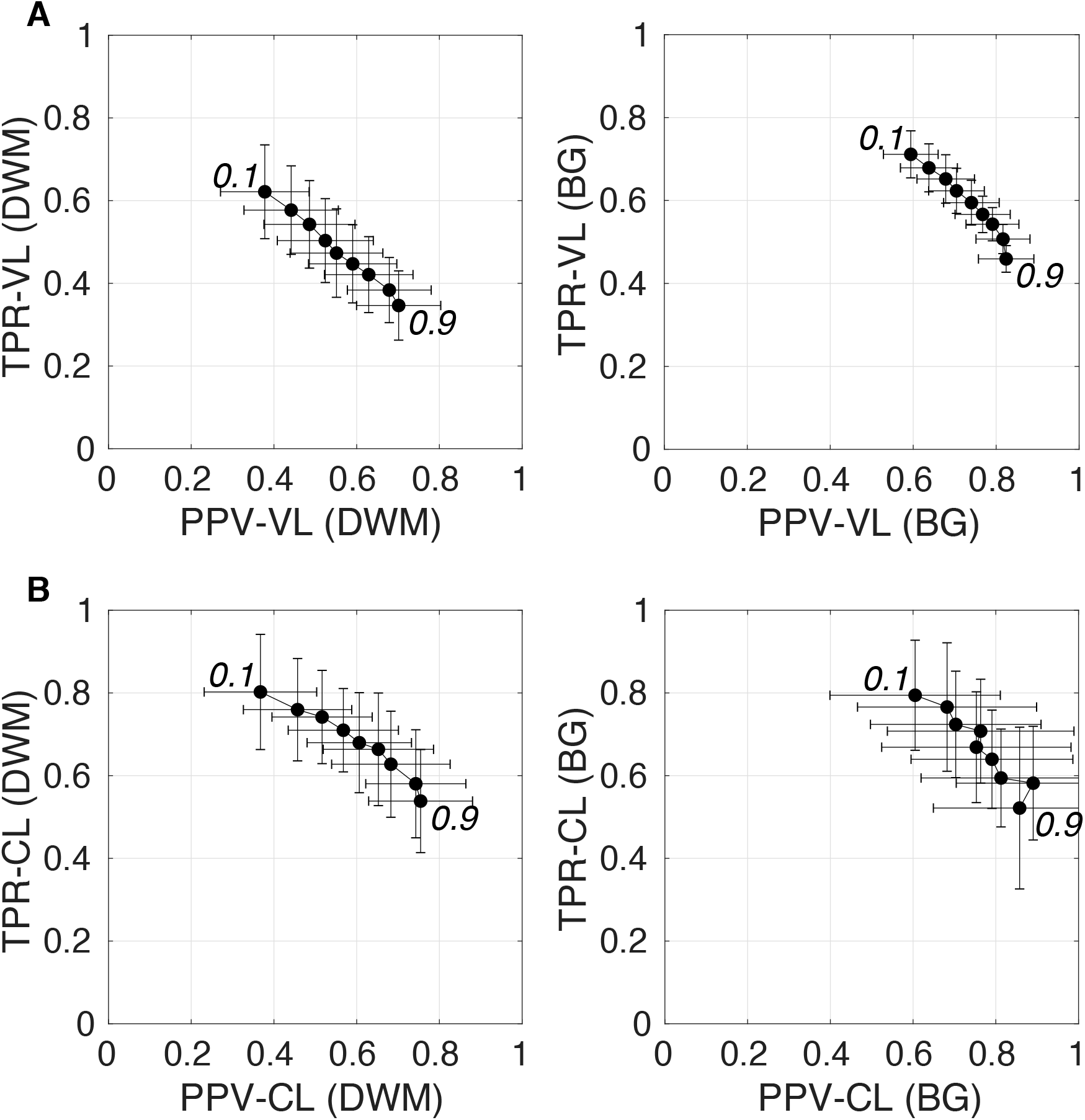
(A) Plot of the true positive rate (TPR) *vs the* positive predictive value (PPV) for the 9 amplitude prediction map thresholds (PredMap-Thrs, 0.1 to 0.9 in steps of 0.1) in the deep white matter (DWM, left) and basal ganglia (BG, right) at the voxel level (VL) (A) and cluster level (CL) (B). Each point gives the average and standard deviation of indexes across the 10 individuals of the EVAL dataset.

**Table 3:** Voxel level (VL) detection true positive rate (TPR), positive predictive value (PPV) and the harmonic mean (Dice) for the 9 amplitude prediction map thresholds (PredMap-Thrs) in deep white matter (DWM) and the basal ganglia (BG).

| Tissue | PredMap-Thr | TPR-VL | PPV-VL | Dice-VL |
| --- | --- | --- | --- | --- |
| **DWM** | 0.1 | 0.62* | 0.38 | 0.47 |
|  | 0.2 | 0.58 | 0.44 | 0.50 |
|  | 0.3 | 0.54 | 0.49 | 0.51 |
|  | 0.4 | 0.50 | 0.52 | 0.51 |
|  | 0.5 | 0.47 | 0.55 | 0.51* |
|  | 0.6 | 0.45 | 0.59 | 0.51 |
|  | 0.7 | 0.42 | 0.63 | 0.50 |
|  | 0.8 | 0.38 | 0.68 | 0.49 |
|  | 0.9 | 0.35 | 0.70* | 0.46 |
| **BG** | 0.1 | 0.71* | 0.59 | 0.65 |
|  | 0.2 | 0.68 | 0.64 | 0.66 |
|  | 0.3 | 0.65 | 0.68 | 0.66 |
|  | 0.4 | 0.62 | 0.70 | 0.66* |
|  | 0.5 | 0.59 | 0.74 | 0.66 |
|  | 0.6 | 0.57 | 0.77 | 0.65 |
|  | 0.7 | 0.54 | 0.79 | 0.64 |
|  | 0.8 | 0.51 | 0.82 | 0.63 |
|  | 0.9 | 0.46 | 0.82* | 0.59 |

\*: Best values

**Table 4:** Cluster level (CL) detection true positive rate (TPR), positive predictive value (PPV) and the harmonic mean (Dice) for the 9 amplitude prediction map thresholds (PredMap-Thrs) in deep white matter (DWM) and the basal ganglia (BG).

| Tissue | PredMap-Thr | TPR-CL | PPV-CL | Dice-CL |
| --- | --- | --- | --- | --- |
| **DWM** | 0.1 | 0.80* | 0.37 | 0.50 |
|  | 0.2 | 0.76 | 0.46 | 0.57 |
|  | 0.3 | 0.74 | 0.52 | 0.61 |
|  | 0.4 | 0.71 | 0.57 | 0.63 |
|  | 0.5 | 0.68 | 0.61 | 0.64 |
|  | 0.6 | 0.66 | 0.65 | 0.66* |
|  | 0.7 | 0.63 | 0.68 | 0.65 |
|  | 0.8 | 0.58 | 0.74 | 0.65 |
|  | 0.9 | 0.54 | 0.75* | 0.63 |
| **BG** | 0.1 | 0.79* | 0.60 | 0.69 |
|  | 0.2 | 0.77 | 0.68 | 0.72 |
|  | 0.3 | 0.72 | 0.70 | 0.71 |
|  | 0.4 | 0.71 | 0.76 | 0.73* |
|  | 0.5 | 0.67 | 0.75 | 0.71 |
|  | 0.6 | 0.64 | 0.79 | 0.71 |
|  | 0.7 | 0.59 | 0.81 | 0.69 |
|  | 0.8 | 0.58 | 0.89* | 0.70 |
|  | 0.9 | 0.52 | 0.86 | 0.65 |

\*: Best values

#### 3.4.2 PVS segmentation model performance for varying PVS cluster sizes

Figure 8 shows grid plots of TPR-CL *versus* PPV-CL (Figure 8.A and 8.B for DWM and BG PVSs, respectively) and surface plots of the Dice-CL (Figure 8.C and 8.C for DWM and BG PVSs, respectively) at different intensity (0.1 to 0.9) and cluster size (0 to 15 voxels) thresholds. The values of each of these indexes are provided in the supplementary material.

**Figure 8.**
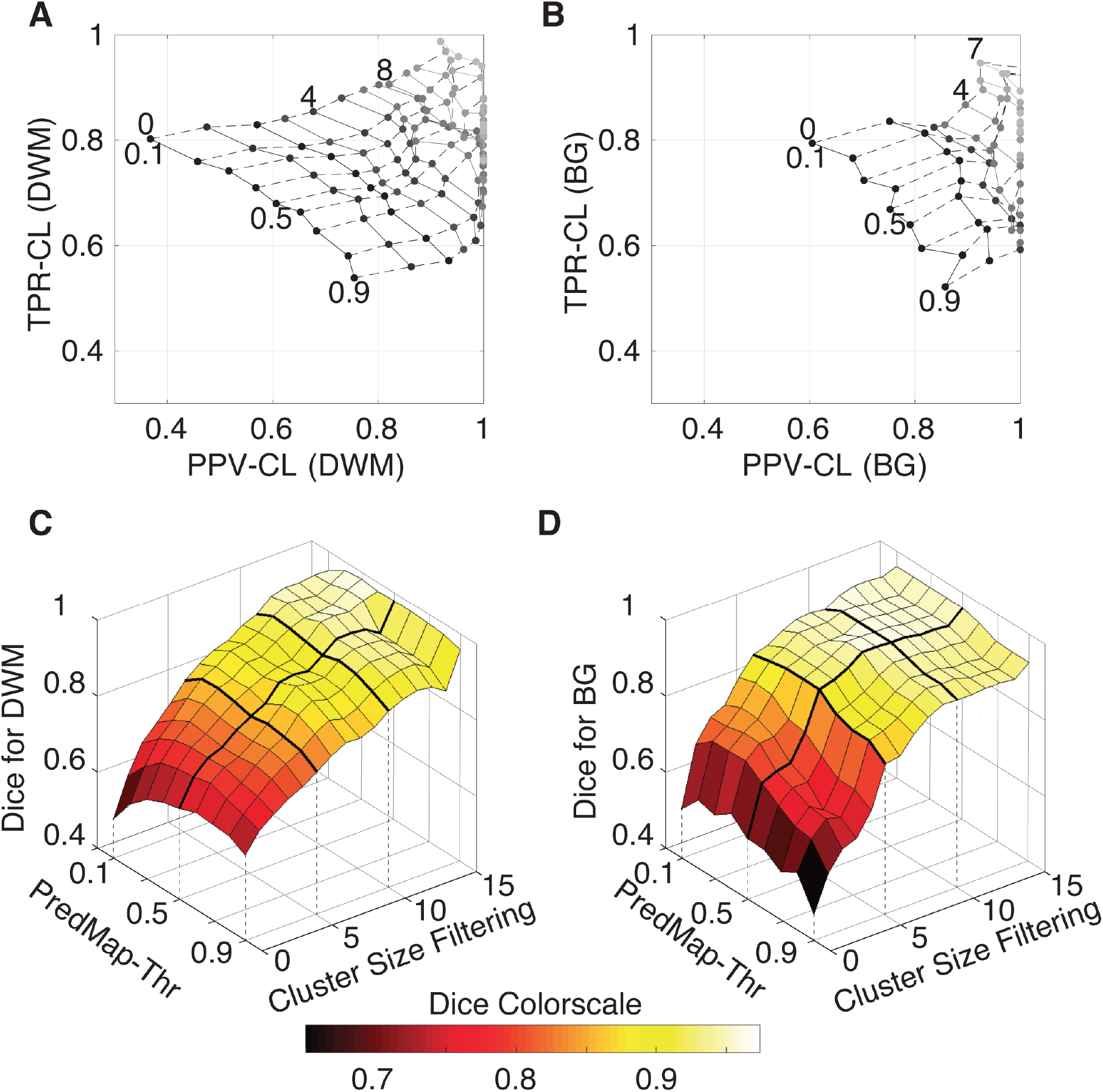
A) Cluster-level true positive rate (TPR-CL) versus the positive predictive value (PPV-CL) of PVS detection in deep white matter (DWM) for the EVAL dataset (N=10). Index values are shown with masking clusters with increasing size (dashed lines, 0 to 15 voxels) and 9 amplitude prediction map thresholds (PredMap-Thrs, solid lines, 0.1 to 0.9 in steps of 0.1). B) Same for detection of basal ganglia (BG) PVS, increasing the cluster size threshold from 0 to 7 voxels. C) Dice scores of PVS detection in deep white matter for the EVAL dataset (N=10) at the 9 PredMap-Thr and masking clusters with increasing size (0 to 15 voxels). D) Same as C but for basal ganglia PVS detection.

As expected, better performance for both indexes was observed when the cluster size threshold was increased. At lower values (1 to 5 voxels), the cluster size threshold had a major impact on PPVs regardless of the intensity threshold. For instance, for DWM PVSs, ignoring clusters made of only one voxel increased the PPV by approximately 15% while increasing TPRs by only a few percentage points. Furthermore, Dice scores above 0.9 could be obtained regardless of the threshold when removing clusters of size 15 or less (Figure 8.C). Data above a cluster threshold of 15 are not shown, as less than half of the EVAL volumes had at least one cluster in the FP or FN classes at this threshold.

For the BG PVSs the grid plot (Figure 8.B) is less smooth due to a lower number of PVSs in the BG (101 clusters on the whole EVAL dataset) compared to the number in DWM (882 clusters). Nevertheless, a marked increase was observed in both the TPR and the PPV after removing the PVS of one voxel. The progression toward optimal values of the PPV was faster in BG PVSs than in DWM PVSs. The Dice index reached 0.9 (regardless of the *PredMap-Thr*) using a cluster threshold size of 7 voxels.

#### 3.4.3 PVS segmentation model performance at predicting PVS cluster sizes

Figure 9 shows the model-predicted PVS size *versus* the size of its corresponding manually annotated PVS. At the *PredMap-Thr* of 0.5, the linear regression slopes were 0.58 (R^2^=0.72) and 0.87 (R^2^=0.93) for the DWM and BG PVSs, respectively. With a 0.1 *PredMap-Thr*, the slopes were closer to 1 (0.83 and 1.2 for DWM and BG PVS, respectively) and were markedly lower with a *PredMap-Thr* of 0.9 (0.39 and 0.63). In both cases, the uncertainty analysis demonstrated that there was no bias in the model regardless of the values.

**Figure 9.**
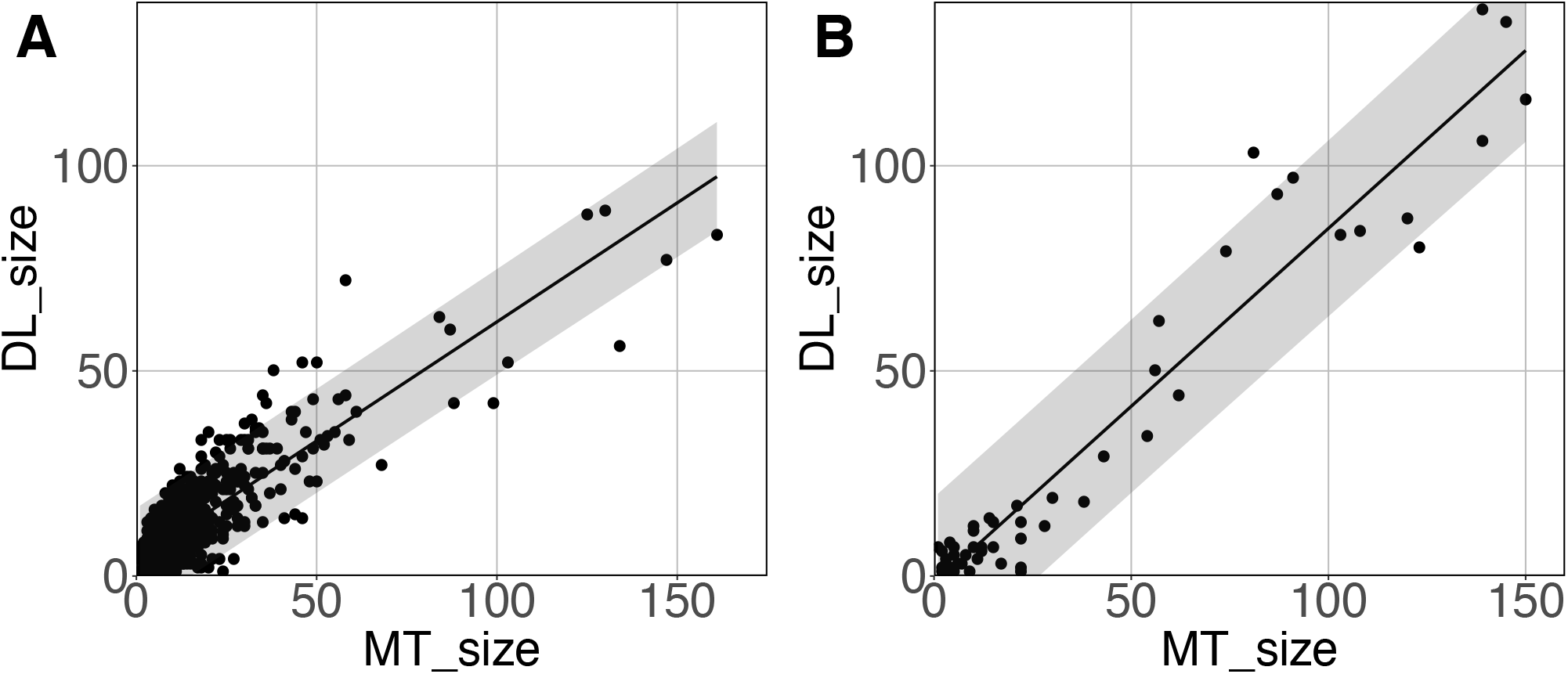
Predicted cluster size (DL_size) versus the manual tracing size (MT_size) for deep white matter (A) and the basal ganglia (B). Both were computed with the amplitude prediction map threshold (PredMap-Thr) of 0.5. Black line: linear regression curve; shaded area: 95% confidence interval of individual values.

### 3.5 Testing the PVS segmentation model of a large subset: ENCOD subset analysis

#### 3.5.1 Relation between the voxel and cluster charge

Figure 10 shows that the relation between the PVS voxel load and the number of clusters can be modeled using a second-order polynomial fit. The adjusted squares of the correlation were 0.94, 0.96 and 0.96 for *PredMap-Thrs* of 0.1, 0.5 and 0.9, respectively.

**Figure 10.**
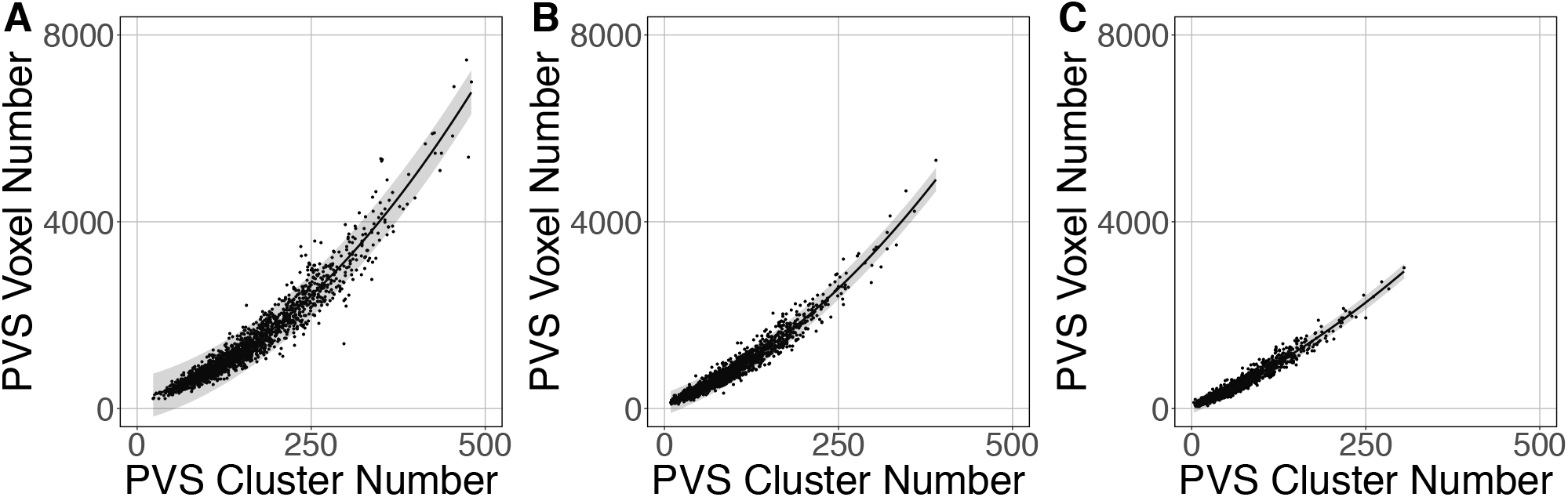
Relationship between the number of clusters and the PVS charge load at 3 amplitude prediction map thresholds (PredMap-Thrs) of 0.1 (A), 0.5 (B) and 0.9 (C). Black line: best second-order polynomial regression curve; shaded area: 95% confidence interval of individual values.

#### 3.5.2 Testing the segmentation model against visual rating results

The logistic regression between the number of DWM PVS clusters (Figure 11.A) and the visual grading rating (Figure 11.B) in the ENCOD set was highly significant for all 3 *PredMap-Thrs* (R^2^_0.5_=0.38, R^2^_0.5_=0.45 see Figure 11.C, R^2^_0.9_=0.47, p <0.001, N=1,782).

**Figure 11.**
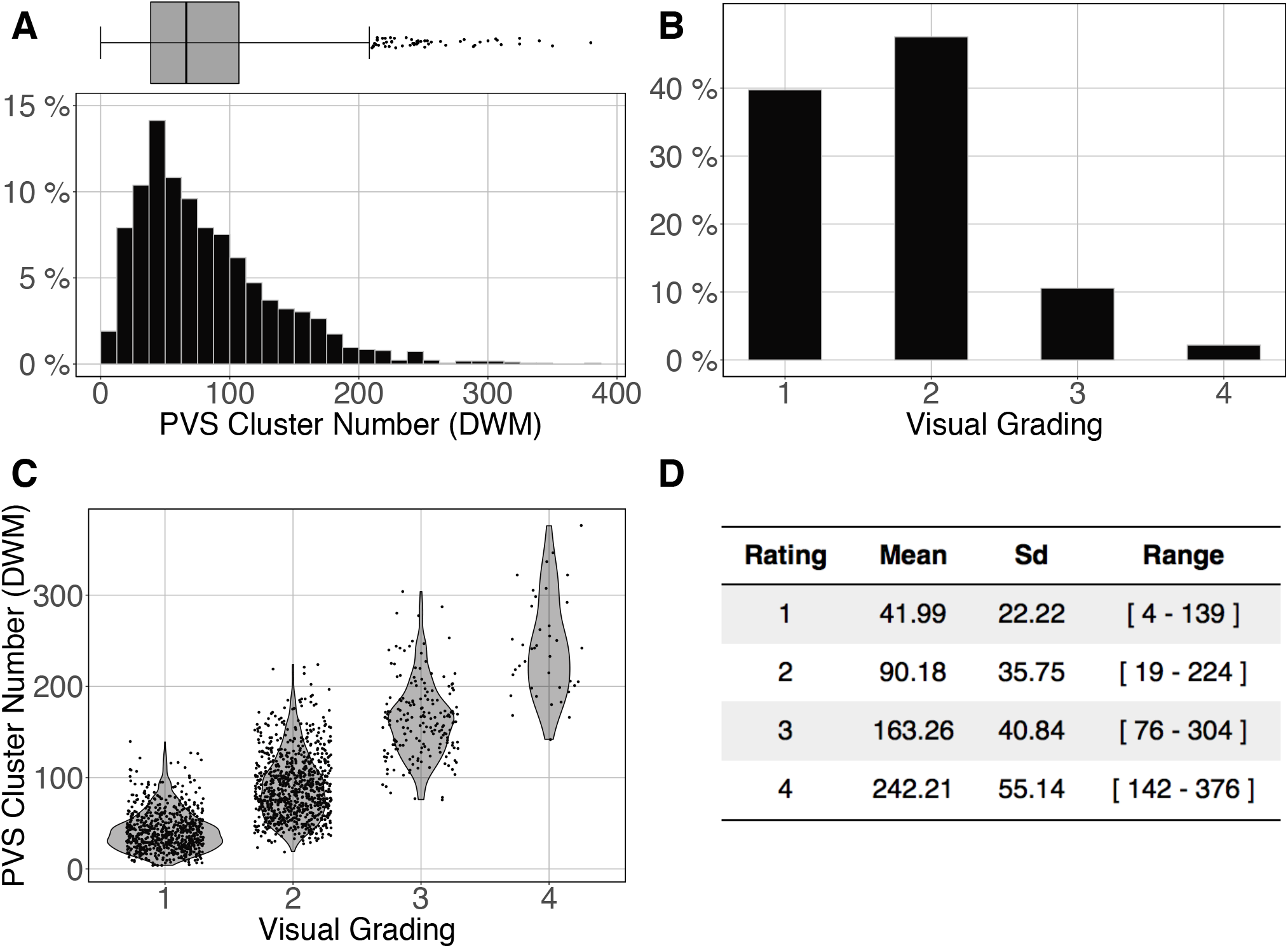
Cluster analysis of the deep white matter (DWM) PVS of the ENCOD set (N=1,782). The prediction maps were thresholded at 0.5. A) Histogram distribution of the number of clusters per subject. B) visual grading. C) Number of clusters versus visual grading (*R*^2^_0.5_=0.45, *p-value* < 10^-4^). D) Summary of PVS load values within the 4 groups.

Figure 12 shows the comparison of the number of BG PVS clusters (Fig. 12.A) and the first 3 levels of visual grading (Figure 12.B) in the ENCOD set. Again the logistic regression showed significant correspondence between the two (R^2^_0.1_=0.02, R^2^_0.5_=0.05 see Figure 11.C, R^2^_0.9_=0.04, p <0.001, N=1782). Note that there were no subjects with BG PVS category of level 4 in the visual rating scale, as such a level is more routinely observed in elderly subjects.

**Figure 12.**
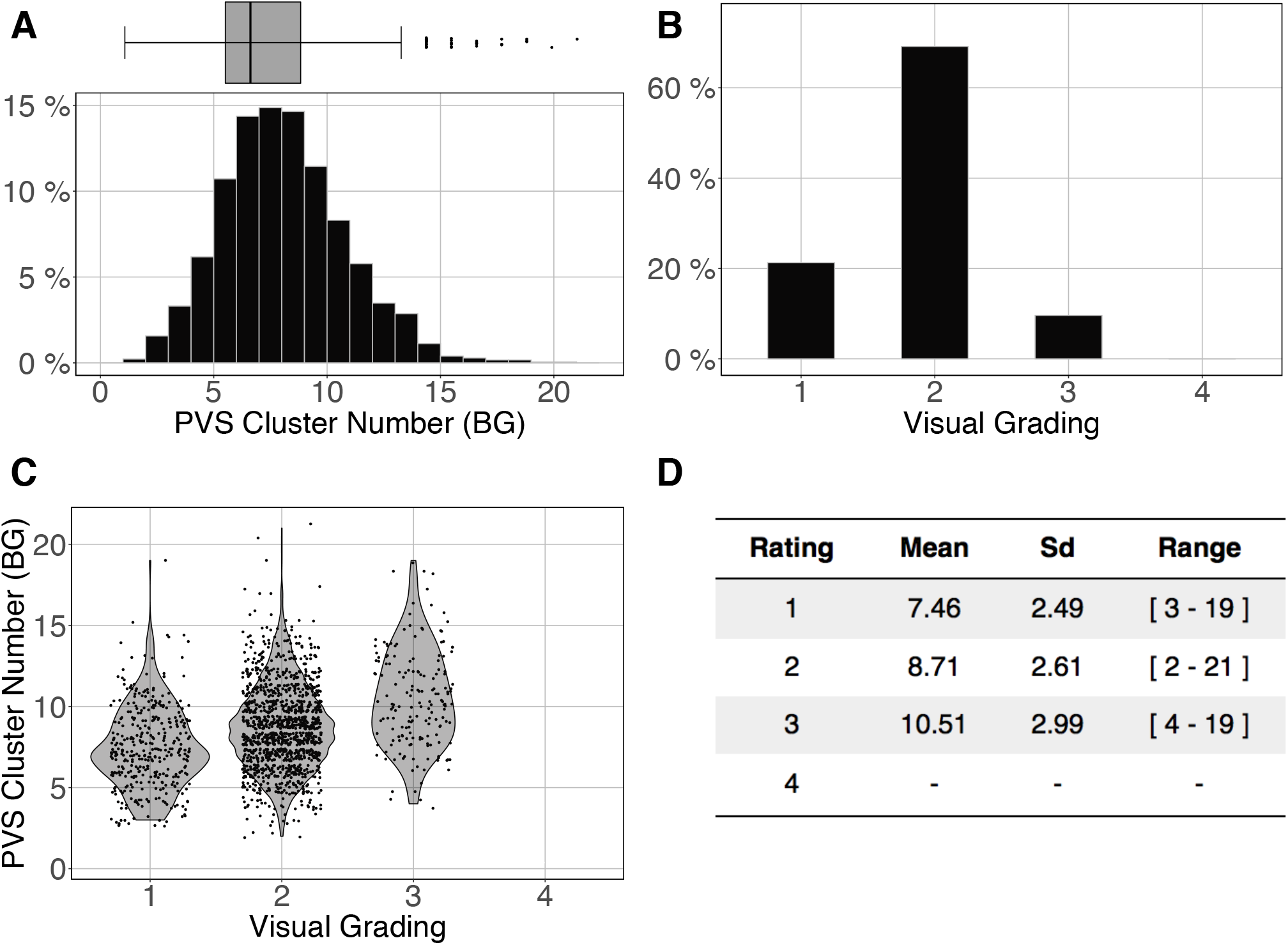
Cluster analysis of the basal ganglia (BG) PVS of the ENCOD set (N=1,782). The prediction maps were thresholded at 0.5. A) Histogram distribution of the number of clusters per subject. B) Visual grading. C) Number of clusters versus visual grading results (*R*^2^_0.5_=0.05, *p-value* < 10^-4^). D) Summary of PVS load values within the 4 groups.

### 3.6 Assessment of the prediction database interoperability

Figure 13.A and B shows the overlap of the number of cluster distributions in the MRi-Share and BIL&GIN datasets. Although the datasets were age-matched, more PVSs were observed in the BIL&GIN subjects when not filtering small clusters (Figure 13.A). Note that this result can also be clearly seen on the cumulative plot of both distributions, also called the QQ plot, presented in Figure 13.C. The difference was quantified by a Kolmogorov-Smirnov test, which showed a significant difference in the distributions (d = 0.23, p-value < 10^-4^). As we showed in the previous sections that filtering out the small cluster improves the reliability of the algorithm, we aimed to find the level of cluster filtering that made the distributions more comparable. After removing clusters below 5 voxels, both distributions (see Figure 13.B and the QQ plot in Figure 13.D) were not different according to the Kolmogorov-Smirnov test (d = 0.077, p-value = 0.058). The discrepancies remained (see Figure 13.D) mainly for the few subjects showing the highest number of PVSs, with more subjects in the MRi-Share dataset having the higher number of PVS clusters than in the BIL&GIN dataset.

**Figure 13:**
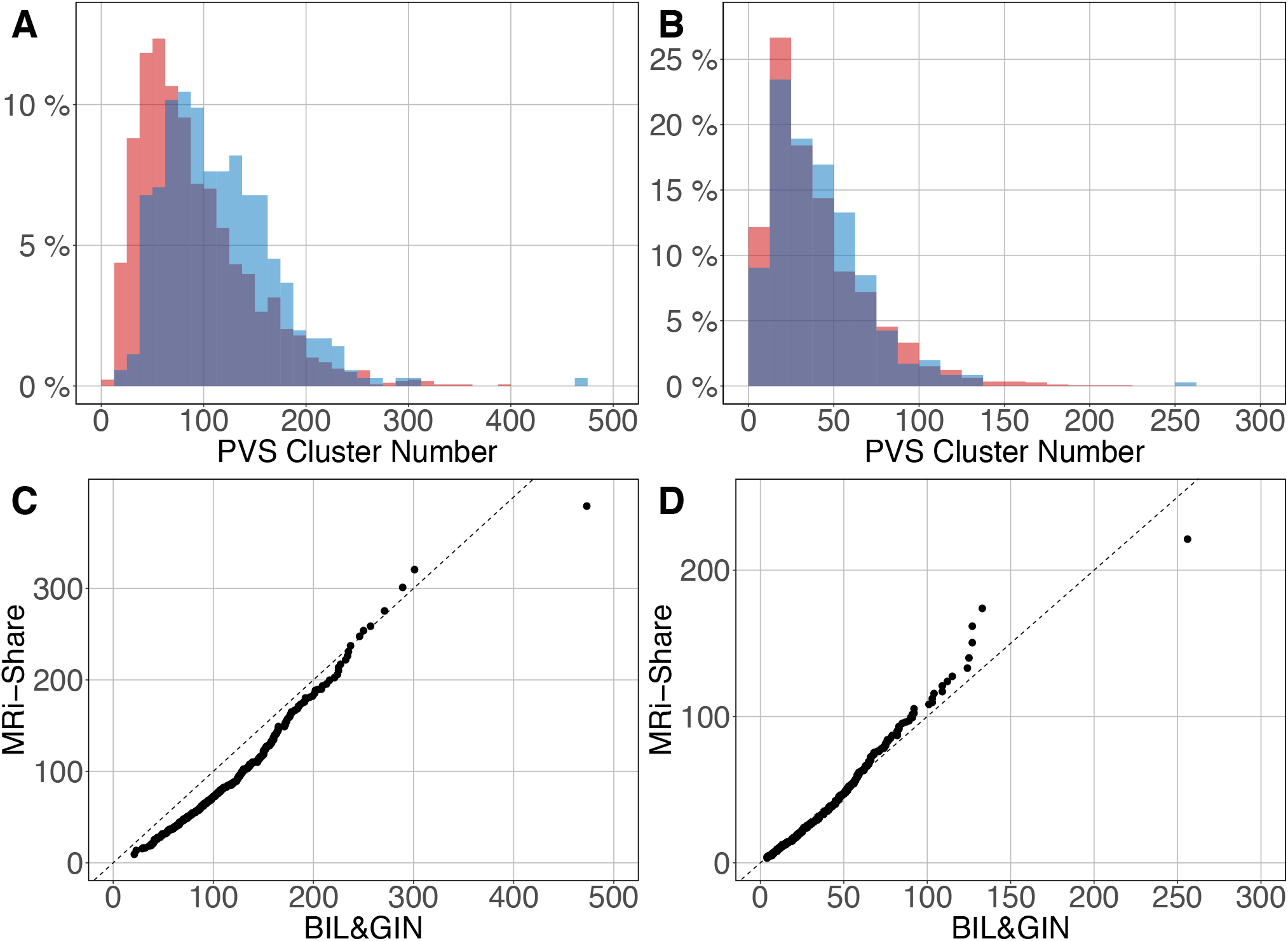
Overlap of the distribution of the number of clusters per subject for MRi-Share (red) and BIL&GIN (blue) cohorts computed at an amplitude threshold of 0.5 and without (A) and with (B) filtering clusters that were <=5 in size. C) and D) QQ plots of the distributions shown in A) and B), respectively.

## 4 Discussion

### 4.1 Methodological issues

#### 4.1.1 Model building

The U-net architecture for our machine learning model was chosen for its segmentation performance with a small training set. We initially tried to implement our algorithm with a training set made of only 10 images, but such a small training set did not allow us to obtain stable and reproducible results; in some cases, the training phase did not converge on a solution or took a very long time (more than 1000 epochs). We then observed that initializing the U-net model with the weights determined by training the autoencoder on the ENCOD set helped provide stable results with fewer epochs (usually under 200 epochs).

The main hyperparameters chosen for the U-net used in our work were based on the U-net topology described in (Ronneberger et al., 2015) adapted for 3D images and constrained by the available RAM in a GPU where the model parameters together with a batch of images should be stored. The initial number of kernels for the first stage of convolutions is important since it provides the basis for the number of features that may be extracted at each level of resolution: the larger the number is, the more features it can extract; however, the resolutionhas a large impact on the size of the model. The number of stages is also important since it adds information for each successive resolution, whereas the number of convolutions for each stage, when greater than 1, seems to be less important. As stated above, 7 stages are needed for the model for going to the bottom stage with an image size of (1, 1, 1), which only allows for an initial number of kernels of 8 with the GPU we used. Since training the model took approximately 12 hours, it was difficult to perform a full grid search or even to use the kind of optimization described in (Bergstra et al., 2013). Obvious tests, such as decreasing the number of stages to increase the number of kernels, were performed without significant gains.

Image cropping during the preprocessing phase provided a 52% reduction in data volume size, which authorized two larger batches and thus increased the training speed. We also tested the segmentation using data registered in MNI stereotaxic space with the same sampling as the acquisition (1 mm^3^); however, both VL TPR and PPV indexes were lower than without normalization (15 and 25%, respectively, for an amplitude PredMap-Thr of 0.5). Visual analysis of the prediction showed that it could be attributed to smoothing due to interpolation. In fact, smoothing small elongated structures such as PVS makes it more difficult for them to be detected because of the induced partial volume effect. We also tested the increase in the training set (both with and without data stereotaxic normalization) using flipping with respect to the interhemispheric plane, but neither case provided any significant increases in the TPR and PPV indexes.

Finally, we tested a more complex U-net topology with U-net++ (Zhou et al., 2020), but it proved to be a failure for the small amplitude PredMap-Thr (0.1 and 0.2) with TPR near 1 and PPV near 0 and marginally better at the other amplitude thresholds for the TPR and worst for the PPV. For the sake of parsimony, we chose to use the more basic U-net topology.

#### 4.1.2 Model reproducibility and robustness

The random nature of the initialization was tested and proved not to be an issue; thus, we did not implement a double-level procedure, such as repeating the training for each fold. This would lengthen the training process (x5) and require the management of 5 times the number of parameters.

The size of the training set proved to be an issue when it was reduced to 20. At this size, using an amplitude threshold below 0.5, the segmentation failed for some of the folds. With a weaker reduction of the training set at 30 data points, the only visible effect was a decrease in the TPR (and not the PPV) of a few percentage points—values that were well below the uncertainties of the measured values (see Figure 7). Such good results were expected when using *U-net* technology. However, to both maintain robustness and limit overlearning, the training dataset size should be as large as possible.

### 4.2 Algorithm performance

We presented our algorithm performance at both the VL and CL. Regardless of the amplitude threshold, both TPR values and PPVs were higher at the CL than at the VL. This discrepancy was related to an imbalance in FNs, possibly due to the small difference in shape between manually traced and predicted clusters. While it was difficult to obtain a definite answer, it is probable that the full extent of the PVS was not included in the manual tracing, leading to the observed discrepancies. Nevertheless, indexes measured at both levels exhibited a strong correlation with each other through a monotonic second-order polynomial relationship. Whatever the chosen index level (voxel or cluster), TPR (resp. PPV) decreased (resp. increased) when the amplitude threshold increased, which led to the best Sorensen-Dice coefficient value with a medium threshold. We proposed that the 0.5 threshold should be used if one does not have any specific reason to favor one of the TPR or PPV indexes.

The algorithm was trained without taking the PVS location into account, namely, whether the PVS was located in the DWM or in the BG. However, for the reasons explained above (see section 1.2), the algorithm predictions on the EVAL and ENCOD sets were analyzed independently for the two locations. From manual tracing, we observed in this dataset of young subjects that the deep WM-located PVS was more numerous and smaller than the BG PVS. Averaging cluster-level indexes across the EVAL dataset demonstrated that, regardless of the amplitude PredMap-Thr, TPR values were equivalent for the 2 locations, whereas PPVs were higher for BG than for DWM. As PPVs are dependent on the FP rate, we investigated their behavior when filtering out clusters according to their size. For both locations, filtering out clusters of 1 voxel size led to a large increase in both index values. This was expected since no PVS of one voxel size was manually traced. Nevertheless, increasing the size of filtered out clusters led to further increases in values of both index values with the increase being larger for PPVs than for TPRs. To summarize, the good performance of our DL algorithm could be improved by considering only PVSs of larger sizes, a feature that could be very interesting when the goal is to detect and quantify dilated PVS.

Likewise, we showed that the PVS sizes were linearly related to the true sizes, albeit underestimated, and the degree of underestimation depended on the chosen amplitude PredMap-Thr. The best estimation of PVS size was obtained with the lowest threshold, albeit at the expense of lowered PPVs and Sorensen-Dice values.

Comparing the PVS prediction by the algorithm with their visual rating on the ENCOD dataset provided important information regarding the clinical utility of the algorithm. Although a visual rating of the PVS burden was made on only one slice for each location (DWM or BG), we observed a very significant agreement between PVS global burdens estimated by the algorithm and by visual rating (p<0.0001) in both locations. For the best results, nearly 50% of the visual rating variance explained by the algorithm predictions was observed for the number of DWM clusters estimated with an amplitude PredMap-Thr above 0.5. Though also significant, the BG PVS explained-variance was 5 times lower, due in part to very unbalanced frequencies of rating for the visual scale degrees; 65% of individuals were rated as degree 3 and none were rated as degree 4. Interoperability of imaging marker detection algorithms is crucial, especially in the context of population neuroimaging in which multiple databases must be jointly analyzed. This can be achieved either by retraining the neural network for each database or by applying the neural network previously trained on one dataset to the other datasets. The latter approach is preferable since the former would require manual tracing of at least 40 subjects for each dataset (30 subjects for the training validation and 10 subjects for the test); manual tracing for this number of subjects is time consuming and could introduce some bias if the operators are not the same. Here, we applied the neural network trained on the MRi-Share dataset to the detection of PVSs in images acquired on a different scanner in a different sample of individuals having the same age range. The PVS number distributions were found to be very similar for the two datasets and not significantly different when removing clusters with sizes less than 5 voxels.

### 4.3 Comparison to other segmentation methodologies

As stated in the introduction, the different segmentation methods proposed in the literature fall into two broad categories: those based mainly on image processing designed to enhance PVS visibility on the image and those that emphasize machine learning classifications and increasingly, DL-based approaches. In fact, this subdivision is not as clear-cut as some methods of the former category often use machine learning classifiers after image enhancement (support vector machine (SVM), (Gonzalez-Castro et al., 2017) or random forests (Zhang et al., 2017)), while some of the latter category used enhanced images as input for the neural networks (Lian et al., 2018). Regardless of the category, the performance is typically evaluated with several different indexes, and the choice of evaluation method is dictated by what is available as the ground truth. Briefly, the TRP (also known as *sensitivity*) and PPV (also known as *precision*) are often reported whenever the ground truth is based on PVS voxel manual tracing, whereas Pearson correlation or Lin’s coefficient is used when only the number of PVSs is available. We computed similar indexes in order to facilitate the comparison of the performance of our algorithm to those in the literature.

When reviewing and comparing existing PVS detection algorithms, several other factors should be taken into account. First is the quality and type of the input image used for the PVS detection, such as the strength of the acquisition MR scanner and image resolution. Some studies used data acquired either at 1.5 T (Dubost et al., 2020; Gonzalez-Castro et al., 2017), 3T ((Ballerini et al., 2018; Boespflug et al., 2018; Sepehrband et al., 2019; Sudre et al., 2019) and our data) or 7T (Jung et al., 2019; Lian et al., 2018). Up to 3T, the data are most commonly acquired with approximately the same 1 mm^3^ sampling size, whereas with 7T scanners, sampling is usually 8 times higher, providing a crucial advantage for detecting small DWM PVS. Notably, the detection of thin features like PVS, especially in DWM, is much improved at high resolution imaging; thus, comparison of the present work with the literature is only meaningful for data acquired on 3T and 1.5T scanners. The imaging sequence is another important factor, as T2-weighted sequences provide better contrast (Zong et al., 2016) for PVS, whereas we worked with T1w images. However, the T1w has the advantage of reduced potential confusion between PVS and WM hyperintensities, which is a common problem when working with T2-weighted images. Another important advantage of an algorithm trained with T1w is the fact that the T1w images are the most commonly available 3D images with millimeter resolution, and it opens the possibility to quantify PVS in datasets that were not originally designed to detect PVS.

Overall, our algorithm exhibited better performance when compared to the performance measures reported in previous studies that used T2-weighted images acquired at 3T or less. In the “image processing” category, Ballerini et al. (Ballerini et al., 2018), using a Frangi filter, reported a Pearson correlation between visual rating and either the total PVS volume burden (r = 0.53) or the total PVS number (r= 0.67). The corresponding values were 0.79 and 0.77, respectively, based on the number of clusters derived from our algorithm (0.5 amplitude prediction map threshold) and the visual rating in the ENCOD set. Using filtering techniques and morphological constraints, Boespflug et al. (Boespflug et al., 2018) reported correlations of 0.58 and 0.76 for the PVS volume and number of clusters; these values indicated worse performance for the total volume burden and equivalent performance for the number of PVS when compared to ours. Finally, using the Frangi filter to create “vesselness” images and nonlocal mean filtering techniques, Sepehrband et al. (Sepehrband et al., 2019) reported a visual rating to cluster number correlation of 0.61 (filtering out the less than 5 voxel clusters), again showing lower correlation than ours. In the 2 studies using a DL methodology, Dubost et al. (Dubost et al., 2020) used a convolutional neural network (CNN) weakly supervised detection approach with attention maps to create class activation maps and reported a sensitivity (called TPR in our study) of 0.53 and 0.60 for DWM and BG PVS, respectively, whereas we reported values of 0.62 and 0.73 for the PVS in the two regions. Sudre et al. (Sudre et al., 2019), using region-based CNN (R-CNN), reported a sensitivity of 0.73 for PVSs above 5 voxels in size, whereas we reported a value of 0.91 under the same conditions (see Figure 9).

Even when comparing with studies using T2-weighted images acquired at 7T, our results indicate competitive performance of our algorithm: Zhang et al. (Zhang et al., 2017), using a structured random field on extracted vascular features, reported a Dice coefficient of 0.66, which was identical to what we obtained for BG PVS, whereas for DWM PVS, it was lower (0.51). However, the slightly better performance in Zhang study for DWM PVS is to be expected, as a higher sampling rate increases the detectability of small PVSs that are mainly located in the DWM. Similarly, Lian et al. (Lian et al., 2018), who used a U-net approach called M2EDN on higher resolution (0.5^3^ mm^3^) T2-weighted images, reported 0.77+-0.04 Dice values (at the VL regardless of the localization) compared to 0.66 and 0.51 for the BG and DWM VRS, respectively, in our study. Without the same resolution in the data while searching small objects, it is difficult to interpret those quantitative differences; thus, we will discuss the differences in the methodology. First, our implementation allows us to process the whole-brain volume simultaneously as opposed to patches of the volume as done in M2EDN. By doing so, we avoid artificially cutting PVSs in 2 or more parts and thus make it possible to compute the number of PVS and not solely the number of PVS voxels. This choice impacts the architecture of the CNN; while Lian et al. (Lian et al., 2018) choose 3 stages with 64 features per level to accommodate the local nature of the patches with PVS, we choose 7 levels with an increasing number of features (from 8 for the first level to 512 at the deepest level) to accommodate both the local and global patterns of PVS repartition in the volume (PVSs are not located everywhere in the volume). A second difference relies on using or not using the multiscale feature, which is equivalent to the U-net++ topology (Zhou et al., 2020) that we tested in the selection of the best topology (see 3.1.1). We did not select it for further analysis because of failures for the small amplitude PredMap-Thr (0.1 and 0.2) and no univocal amelioration of the evaluation indexes at the other thresholds. Note that Lian et (Lian et al., 2018) did not report such failure, but they also did not report the amplitude PredMap-Thr used to compute their algorithm evaluation indexes. Finally, it must be emphasized that M2EDN is based on multichannel MRI data, including raw T2-weighted images, enhanced T2-weighted images and probability maps. Using the probability map in what Lian et al. (Lian et al., 2018) called “autocontext” did not demonstrate a decisive advantage, and the procedure was limited to one iteration. In addition, we believe that having a portable PVS detection algorithm operating on 3D-T1w images only constitutes an important advantage in favor of our approach since this algorithm could be easily applied to most of the existing MRI databases and/or clinical brain MRI protocols that often include a high-resolution 3D T1w acquisition.

## 5 Conclusion

We implemented a U-net-based DL algorithm for the 3D detection of PVS on T1w images both in the DWM and the BG area. Overall, when considering images of comparable resolution, our U-net-based DL PVS segmentation algorithm exhibited better performance than that of previously published methods working with T2-weighted images, whether based on signal processing or DL methods. The algorithm performance and its interoperability for 3T T1w data are important features in the context of both routine clinical analysis and mega- or meta-analysis of PVS across databases, as 3D millimeter T1 images are available for many existing neuroimaging databases.

## 6 Conflict of Interest

The authors declare that the research was conducted in the absence of any commercial or financial relationships that could be construed as a potential conflict of interest.

## 7 Funding

This work has been supported by a grant from “La Fondation pour la Recherche Médicale” (DIC202161236446 WAIMEA, B Mazoyer PI, PLB, AL, LL, AT) and by a grant overseen by the French National Research Agency (ANR) as part of the “Investissements d’Avenir” Program ANR-18-RHUS-002. This work was supported by a grant from the French National Research Agency (ANR-16-LCV2-0006-01, LABCOM Ginesislab M Joliot and P Boutinaud PIs, ZH, VN, VV). The i-Share study has received funding from the ANR (Agence Nationale de la Recherche) via the ‘Investissements d’Avenir’ programme (grant ANR-10-COHO-05, C Tzourio PI). Supplementary funding was received from the Conseil Régional of Nouvelle-Aquitaine (ref. 4370420). The MRi-Share cohort was supported by grant ANR-10-LABX-57 (B Mazoyer PI). The work was also supported by the “France Investissements d’Avenir” program (ANR–10–IDEX-03-0, C Tzourio PI).

## 8 Acknowledgments

We thank Pierre-Louis Bazin for providing the Medical Image Processing, Analysis and Visualization software.

## 9 Author contributions

**Philippe Boutinaud**: Methodology, Software, Validation, Writing - Original Draft, Writing - Review & Editing, Supervision, Project administration, Funding acquisition. **Ami Tsuchida**: Data Curation, Writing - Original Draft, Writing - Review & Editing. **Alexandre Laurent**: Software, Writing - Original Draft, Validation, Writing - Review & Editing, Visualization. **Filipa Adonias**: Validation. **Zara Hanifelo**: Software. **Victor Nozais**: Software, Writing - Original Draft, Writing - Review & Editing. **Violaine Verrecchia**: Software, Writing - Original Draft, Writing - Review & Editing. **Leonie Lampe**: Data Curation. **Junyi Zhang**: Data Curation. **Yi-Cheng Zhu**: Data Curation. **Christophe Tzourio**: Investigation, Writing - Review & Editing, Funding acquisition. **Bernard Mazoyer**: Conceptualization, Formal analysis, Investigation, Writing - Original Draft, Writing - Review & Editing, Supervision, Project administration, Funding acquisition. **Marc Joliot**: Conceptualization, Formal analysis, Investigation, Writing - Original Draft, Writing - Review & Editing, Visualization, Supervision, Project administration, Funding acquisition.

## 11 Supplementary Material

“Cluster_amp_size.xls file”: TPR, PPV, and Dice-Sorensen indexes for PVSs located in the DWM and the BG are provided at 9 amplitude thresholds of the prediction map threshold (PredMap-Thr, 0.1 to 0.9 in steps of 0.1) and cluster size thresholds (f0: no filtering, f15: clusters with volume greater than 15mm^3^).

## Notes

### Competing Interest Statement

The authors have declared no competing interest.

